# Exome sequencing and genotyping identify a rare variant in *NLRP7* gene associated with ulcerative colitis

**DOI:** 10.1101/182113

**Authors:** Alexandros Onoufriadis, Kristina Stone, Antreas Katsiamides, Ariella Amar, Yasmin Omar, Katrina de Lange, Kirstin Taylor, Jeffrey C. Barrett, Richard Pollok, Bu’Hussain Hayee, John C. Mansfield, Jeremy D. Sanderson, Michael A. Simpson, Christopher G. Mathew, Natalie J. Prescott

## Abstract

**Background and aims:** Although genome-wide association studies (GWAS) in inflammatory bowel disease (IBD) have identified a large number of common disease susceptibility alleles for both Crohn’s disease (CD) and ulcerative colitis (UC), a substantial fraction of IBD heritability remains unexplained, suggesting that rare coding genetic variants may also have a role in pathogenesis. We used high-throughput sequencing in families with multiple cases of IBD, followed by genotyping of cases and controls, to investigate whether rare protein altering genetic variants are associated with susceptibility to IBD.

**Methods:** Whole exome sequencing was carried out in 10 families in which 3 or more individuals were affected with IBD. A stepwise filtering approach was applied to exome variants to identify potential causal variants. Follow-up genotyping was performed in 6,025 IBD cases (2,948 CD; 3,077 UC) and 7,238 controls.

**Results:** Our exome variant analysis revealed coding variants in the *NLRP7* gene that were present in affected individuals in two distinct families. Genotyping of the two variants, p.S361L and p.R801H, in IBD cases and controls showed that the p.S361L variant was significantly associated with an increased risk of ulcerative colitis (odds ratio 4.79, p=0.0039) and IBD (odds ratio 3.17, p=0.037). A combined analysis of both variants showed suggestive association with an increased risk of IBD (odds ratio 2.77, p=0.018).

**Conclusions:** The results suggest that *NLRP7* signalling and inflammasome formation may be a significant component in the pathogenesis of IBD.

## Introduction

Genome-wide association scans (GWAS) have been very successful in the identification of susceptibility genes for many common, complex disorders [https://www.ebi.ac.uk/gwas/]. GWAS in both major forms of inflammatory bowel disease (IBD), Crohn’s disease (CD) and ulcerative colitis (UC), have been amongst the most productive, and provided a better understanding of the biology of these diseases. There are now around 240 robust genetic associations that have been identified ^1-3^ in IBD. However, most associations either span genomic regions that encompass multiple potential candidate genes or lie within non-coding or gene-poor regions. Their biological significance is therefore often unclear, and they explain only a modest proportion of the estimated heritability (or variation in genetic liability) of IBD. A recent study by the International IBD Genetics Consortium used a fine-mapping approach in 67,852 individuals in an attempt to pinpoint the true disease causing DNA variants at 94 of the top IBD loci ^4^. This identified 18 single independent causal DNA variants at 14 loci with >95% certainty and 27 single independent genetic variants at 26 loci with >50% certainty. Taken together these 45 independent associations at 37 loci were enriched for 13 variants which elicit changes specifically in the protein coding sequences of 7 genes.

It is thought that some of the hidden heritability in IBD may be explained by the association of rare protein-altering variants that are likely to confer a larger risk of disease. It is unlikely that such variants would be detected by the GWAS approach alone, which is designed to identify association with common variants. In recent years, advances in DNA sequencing technology have facilitated gene sequencing on a much larger scale than could previously be attempted, enabling large-scale studies that have led to discovery of complex disease causing variants that previously eluded GWAS studies ^5^. Moreover, studies using targeted high throughput sequencing of genes in IBD GWAS regions have, to date, detected around 30 multiple low frequency variants associated with adult IBD ^6-10^ in 17 genes including multiple variants in the *NOD2* and *IL23R* genes.

A recent UK IBD Genetics Consortium study has undertaken low depth sequencing of the entire genome in 4,280 IBD patients and found only one new low frequency protein-coding (missense) variant in the *ADCY7* gene that increases risk for UC. However, this study also detected evidence for an increased ‘mutational-load’ of rare damaging missense variants in known CD risk genes, although it was estimated that these low frequency variants only accounted for a very small fraction of the variation in genetic liability (heritability) of IBD (<2%) in the general population ^11^. In general, the majority of genetic variants associated with common, complex disorders thus far do not lie within the coding regions of genes. However, this is not the case for some subsets of common disorders with a strong familial predisposition, such as breast cancer and Alzheimer’s disease, in which rare, highly penetrant mutations have a causal role.

In this study we hypothesised families with large numbers of individuals affected with adult-onset IBD are more likely to have resulted from the action of rare or low frequency damaging missense changes in one or a few genes. Such disease causing variants would be easier to find by high depth sequencing of coding genes in individuals from these families, and, if present in genes from known IBD risk loci, might help to identify the causal genes in regions intractable to fine-mapping approaches.

## MATERIALS AND METHODS

### Subjects

A large collection of white British IBD families with three or more affected individuals were recruited with informed consent and institutional ethical approval as described previously ^2^. Genomic DNA was prepared from 10mls of whole blood using the salt/chloroform method described elsewhere^12^. Of these, 10 families were selected for sequencing because they had >3 affected individuals across multiple generations. Overall, 7 of the selected families were affected with only CD, 2 with only UC, and 1 family had both CD and UC affected individuals. DNA samples from a total of 6,025 IBD cases (2,948 CD; 3,077 UC) and 7,238 controls of white British descent described previously ^9^ were used for follow up genotyping and association studies as well as genotype data 16,267 UK IBD cases and 18,843 UK population controls from a recent whole genome sequencing study ^11^.

### Exome Sequencing analysis

Whole exome sequencing (WES) was carried out on 27 affected individuals from 10 families, that is, 2, 3 or 4 individuals from each family. Our sequencing strategy was devised so that, where possible, only the most distantly related and affected family members from each branch of each family were sequenced. Using this strategy, we assumed that rare or low frequency variants that were shared between the exome sequenced affected relatives in a family were highly likely to be identical by descent, therefore optimising the information gained against the cost of WES all the affected individuals. Three micrograms of genomic DNA was sheared to a mean fragment size of 150 bp (Covaris), and the fragments used for Illumina paired-end DNA library preparation and enrichment for target sequences (SureSelect Human All Exon 50Mb kit, Agilent). Enriched DNA fragments were sequenced with 100 bp paired-end reads (GAIIx or HiSeq2000 platform,Illumina). Sequencing reads were aligned to the reference human genome sequence (hg19) using the Novoalign software (Novocraft Technologies). Duplicate and multiply mapping reads were excluded, and the depth and breadth of sequence coverage were calculated using custom scripts and the BedTools package. Single-nucleotide substitutions and small indels were identified with SAMtools, annotated with the ANNOVAR software, and variants called as previously described ^13^. On average, 7 gigabases of sequence were generated per sample; >82% of the target exome was present at >20-fold coverage, and >94% was present at >5-fold coverage. Identified variants were prioritised for follow-up based on the following filters:

i. We assumed a dominant model of inheritance and therefore selected only those variants that were heterozygous,
ii. we looked for variants most likely to have a functional affect/be protein-altering, so we focused on nonsynonymous (missense, nonsense or canonical splice site substitutions), and variants that were absent or present with a frequency of < 1% in the Exome Aggregation Consortium database (ExAc, http://exac.broadinstitute.org/about) or whole exome sequence data from >1000 in house non-IBD individuals.
iii. We selected only those variants that were present in all WES members from each family, using the strategy described above,
iv. Finally, to increase the likelihood of selecting disease causing variants, we selected only those genes in which we had identified one or more variants fulfilling criteria i-iii above in at least two different IBD families.

### Pathogenicity Prediction

A combination of three software tools; SIFT (http://sift.jcvi.org/), PolyPhen2 (http://genetics.bwh.harvard.edu/pph2/) and CADD ^14^ (http://cadd.gs.washington.edu/) were applied to evaluate the pathogenicity of the 34 variants identified through our WES and filtering strategy.

### Sanger sequencing confirmation of *NLRP7* variants

The presence or absence of the rare *NLRP7* variants detected by exome-sequencing was confirmed by Sanger sequencing in the same individuals as well as in additional available family members who were either affected (2 for family GS13, 1 for family GS64) or unaffected (7 for family GS13, 5 for family GS64) to test for co-segregation with IBD. Primers were designed using Primer3 software (http://primer3.sourceforge.net/). PCR products were purified with ExoSAP-IT (GE Healthcare), and sequenced using BigDye Terminator v3.1 chemistry (Life Technologies) on a 3730xl DNA analyser (Life Technologies). Sequence traces were visualised using the Sequencher 5.0 software (Gene Codes) and variants were detected by manual inspection of chromatograms.

### Genotyping

Owing to sequence homology of the coding regions of *NLRP7* with its nearby paralogue *NLRP2*, the two rare variants in *NLRP7*, rs143169084 (p.S361L) and rs140797839 (p.R801H), detected by WES in IBD families were genotyped using primers specifically designed to differentiate both paralogues (sequences available on request). Genotyping assays were performed using the KASP chemistry at LGC Genomics (Hoddesdon, Herts, UK) in a further unrelated cohort of 6,025 IBD cases (2,948 CD and 3,077 UC) and 7,238 population controls to test for genetic association with IBD. Samples that failed to genotype for both SNPs were removed (102 CD, 132 UC and 169 controls). Genotype clusters were visualised using SNPviewer (LGC genomics). Carriage of the minor allele was confirmed by direct Sanger sequencing (as described above) in all identified IBD and control individuals within the genotyping cohort for p.S361L (n=18) and p.R801H (n=5). Asymptotic allelic case-control test for association were performed using PLINK (http://pngu.mgh.harvard.edu/∼purcell/plink/). Association p values were corrected for multiple testing based on Bonferroni correction for 6 tests (2 variants x 3 phenotypes).

The remaining 32 variants detected by WES and variant filtering were assessed for association with IBD using data from a recent UK whole genome sequencing study and large GWAS meta-analysis of 16,267 UK IBD cases and 18,843 UK population controls^11^.

## RESULTS

We conducted a screen for rare, potentially disease causing coding variants by whole-exome sequencing (WES) a subset of the most distantly related affected individuals from ten IBD families. The large number of coding variants that were identified across all individuals was prioritised for follow up by applying a series of stepwise variant filters. Firstly, within each family, we retained only those variants that were heterozygous, present in all WES affected family members and were protein altering, i.e. those that introduce a missense, nonsense, or frameshift change in the protein structure or altered a canonical splice site, which left 21,127 variants. Next, we removed all common variants with an allele frequency reported to be equal to or greater than 1%. This filtering strategy resulted in 694 variants identified in 627 genes including 88 novel variants. Our final filter was to restrict the list of variants for follow up to include only those genes that contained rare variants o-segregating with IBD in WES individuals in two or more IBD families (Materials and Methods), thus providing more than one source of evidence for the implicated gene. This resulted in 34, low frequency, protein altering variants in 17 genes responsible for a range of cellular functions including cell migration, metabolism, transcriptional activation and the innate immune response (Supplementary table 1). One of these genes, *NLRP7*, had a reported functional connection with immune-mediated disease and resided within one of the 240 mapped genetic regions associated with IBD ^1-3^ (chromosome 19 at 55.4Mb, hg19). Our exome data had identified two missense variants within the *NLRP7* gene, p.S361L (c.1082C>T, rs143169084) in family GS13 and p.R801H (c.2402G>A, rs140797839) in family GS64. The presence of these variants was validated by Sanger sequencing in the discovery cases, and further genotyped by Sanger sequencing in all available affected and unaffected members in the two families (Figure 1A). In family GS13, p.S361L was present in all three affected family members with CD and in two unaffected individuals, and was absent in three unaffected relatives and an unrelated spouse who, co-incidentally also has CD. This suggests that if p.S361L is the causal variant in this family it is showing incomplete penetrance. In family GS64, the .R801H variant was present in three of four affected relatives (two with CD; one with UC) and in one unaffected family member. The absence of p.R801H in one affected relative (Figure 1A, II:2) suggests that one or more other variants may have contributed to the development of IBD in this individual.

**Figure 1:**
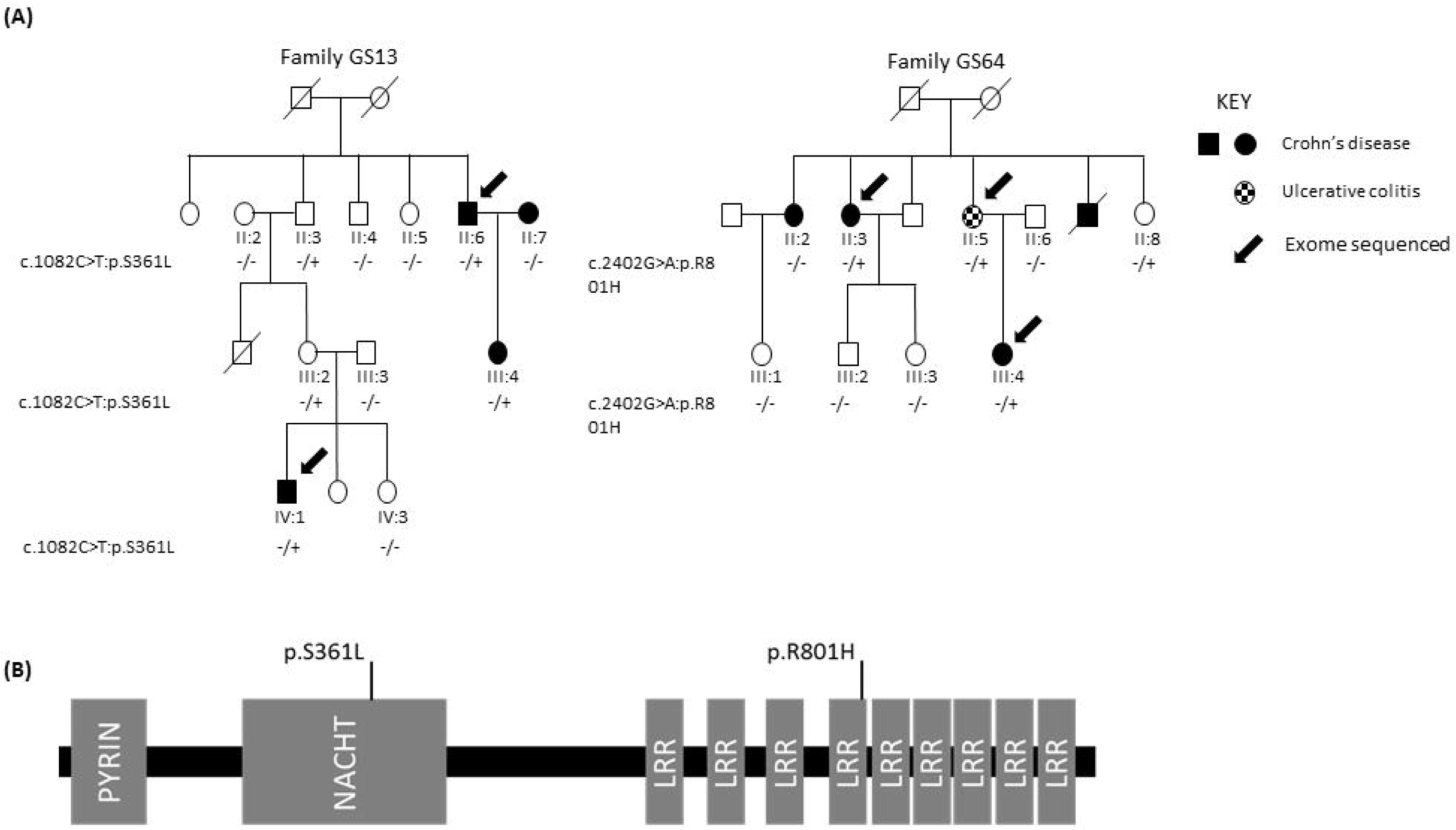
(A) Segregation of the c.1082C>T (p.S361L) variant in pedigree GS13 and c.2402G>A (p.R801H) variant in pedigree GS64. (B) Location of the two protein altering variants with respect to the functional domains of *NLRP7*; Pyrin, NACHT and Leucine-rich repeats (LRR).

The two low frequency *NLRP7* variants were next followed-up by genotyping in a large panel of 6,025 unrelated IBD cases (excluding individuals from the families that underwent WES) and 7,238 population controls to test for association with IBD. Both variants had a high call rate (>99%) and were in Hardy-Weinberg equilibrium in both cases and controls. The minor A allele that encodes the variant leucine residue of *NLRP7* p.S361L had a higher frequency in all IBD subgroups compared to controls (Freq_[CD]_=0.100%,Freq_[UC]_ =0.200%, Freq_[cont]_ =0.035%) and was significantly associated with UC (p=3.9x10^-3^, OR=4.79, 95%CI: 1.60-14.00), also after correction for multiple testing (Table 1). The minor T allele that encodes the variant histidine residue of *NLRP7* p.R801H also had a higher frequency in IBD cases compared to controls but failed to achieve significance on its own (Table 1). Owing to the low frequency of both of the variant alleles in the population we performed a combined analysis of their cumulative frequency in IBD cases compared to controls, and this was significantly different, although not after correction for multiple testing p=0.018, p_corrected_=0.054 (OR=2.77, 95%CI: 1.14-6.75) (Table 1).

**TABLE 1:** Single and combined association analysis of the two protein altering variants in the *NLRP7* gene in 5,801 inflammatory bowel disease cases and 7,074 population controls.

| **p.S361L** | **GG** | **GA** | **AA** | **n** | **MAF** | **P** | **P<sub>corrected</sub>** | **Odds ratio** | **95%CI** |
| --- | --- | --- | --- | --- | --- | --- | --- | --- | --- |
| **CD** | 2846 | 3 | 0 | 2849 | 0.100% | 0.8700 | n.s. | 1.49 | 0.35 - 6.20 |
| **UC** | 2942 | 10 | 0 | 2952 | 0.200% | 0.0039 | 0.0234 | 4.79 | 1.60 - 14.00 |
| **IBD** | 5788 | 13 | 0 | 5801 | 0.110% | 0.0370 | n.s. | 3.17 | 1.13 - 8.90 |
| **Control** | 7069 | 5 | 0 | 7074 | 0.035% | - |  | - | - |

| **p.R801H** | **CC** | **CT** | **TT** | **n** | **MAF** | **P** | **P<sub>corrected</sub>** | **Odds ratio** | **95%CI** |
| --- | --- | --- | --- | --- | --- | --- | --- | --- | --- |
| **CD** | 2804 | 2 | 0 | 2806 | 0.036% | 0.6900 | n.s. | 2.40 | 0.35 - 17.68 |
| **UC** | 2945 | 1 | 0 | 2946 | 0.017% | 1.0000 | n.s. | 1.18 | 0.10 - 13.08 |
| **IBD** | 5749 | 3 | 0 | 5752 | 0.026% | 0.8000 | n.s. | 1.80 | 0.30 -10.90 |
| **Control** | 6984 | 2 | 0 | 6986 | 0.014% | - |  | - | - |

| **combined** | **1/1** | **1/2** | **2/2** | **n** | **Frequency** | **P** | **P<sub>corrected</sub>** | **Odds ratio** | **95%CI** |
| --- | --- | --- | --- | --- | --- | --- | --- | --- | --- |
| **CD** | 2801 | 5 | 0 | 2806 | 0.18% | n.s |  | 1.77 | 0.56 - 5.6 |
| **UC** | 2935 | 11 | 0 | 2946 | 0.37% | 0.003 | 0.009 | 3.73 | 1.44 - 9.63 |
| **IBD** | 5736 | 16 | 0 | 5752 | 0.28% | 0.018 | 0.054 | 2.77 | 1.14 - 6.75 |
| **Control** | 6979 | 7 | 0 | 6986 | 0.10% | - |  | - | - |

The variant p.S361L maps to the NACHT domain of the *NLRP7* protein whilst the p.R801H missense substitution is located in the leucine rich repeat domain (Figure 1B). The p.S361L variant was predicted to be “deleterious” by the CADD pathogenicity software tool (CADD score 23.4), “damaging” by SIFT and “probably damaging” by PolyPhen-2. The p.R801H variant was predicted to be “tolerated” (CADD score 13.99) “neutral” by SIFT and “benign” by PolyPhen-2.

The remaining 32 variants in 16 genes that were identified by our WES and filtering strategy (Supplementary table 1) were followed up by examination of their frequency distribution in a recent large GWAS meta-analysis of 16,267 UK IBD cases and 18,843 UK population ontrols^11^. Of the ten variants that were detected in the GWAS meta-analysis study and passed quality control, two showed evidence of association with IBD (p< 2x10^-3^). These were both in the *TRIM31* gene (supplementary table 1). *TRIM31* is located on chromosome 6p within the human MHC region. One of the associated variants, rs62624473, is a splice site substitution, and the minor allele (freq = 0.9%) is associated with a decreased risk for UC and IBD (p = 0.00174 and 0.00172 respectively; OR = 0.55 and 0.86 respectively) in the GWAS-meta analysis data. This is in contrast to the observed co-segregation of the rare allele with UC in family 107, suggesting this variant is unlikely to be causal of disease. The second variant in *TRIM31* is the missense change p.C48R, rs140451451. This variant is very rare (0.07%), but shows evidence of association with UC and IBD in the GWAS meta-analysis data (p = 2.82x10^-05^ and 5.81x10^-04^ respectively). It has a low CADD score (5.416), and is predicted to be “possibly damaging” and “tolerated” by SIFT and Polyphen2 respectively. Also, we cannot exclude the possibility its association with IBD is a result of its correlation with common variants at the nearby HLA-DRB1 gene in the human MHC region, where there is known to be extensive linkage disequilibrium and multiple IBD associated haplotypes^15^.

## DISCUSSION

Whole exome sequencing of affected individuals in IBD families identified 17 genes in which rare coding variants were present in more than one family. None of these genes have thus far been found to be mutated in monogenic disorders associated with intestinal inflammation ^16^, and only one, *NLRP7*, is located in one of the 240 loci known to be associated with IBD. This suggests that rare, highly penetrant, coding variants in genes from GWAS loci may not play a substantial role in the UK IBD families we have sequenced. The discovery of at least one significant association of a low frequency coding variant in *NLRP7* (p.S361L) with IBD suggests that this and potentially other variants in this gene may predispose individuals to IBD. This is further supported by the prediction that p.S361L would damage the function of the *NLRP7* protein. The *NLRP7* variant p.R801H may be an incidental finding and not causally related to IBD in family GS64. However the co-segregation of the variant allele with three of four individuals with the IBD phenotype in this family and the association of both *NLRP7* variants with IBD in the combined analysis warrants further exploration of these and additional *NLRP7* variants in very large case-control collections and additional IBD families.

The *NLRP7* gene is located on chromosome 19 at position chr19:55,434,877-55,458,873 (hg19) within a known IBD locus. The region was initially identified by GWAS ^1^ to be significantly associated with IBD p=6.5x10^-11^, and spans multiple genes including *NLRP7*, *NLRP2*, KIR2DL1, LILRB4 and 15 others. The top associated SNP in the region is rs11672983, although the two missense variants identified in this study are not in linkage disequilibrium with this SNP (r^2^=0). This locus was not included in the recent fine-mapping study by the International IBD consortium^4^.

Whilst the *NLRP7* variants were identified in families that contained individuals affected by both CD and UC or CD alone, we were only able to detect a significant association signal for p.S361L in UC. However, these findings do not preclude a role for *NLRP7* in CD. It is possible that the lack of association in this study may be because the effect size is smaller in CD (OR = 1.49) and thus lacked sufficient power to detect the smaller effect.

It has been reported that the p.S361L variant is more prevalent in the Ashkenazi Jewish (AJ) population (MAF=0.0438) compared to white European populations (MAF=0.0004) by the gnomAD database (http://gnomad.broadinstitute.org/). This is of interest as IBD is more prevalent in the AJ population^17-19^; however there has been no reported association of *NLRP7* or this region of chromosome 19 with IBD in this population to date^20^, and none of the IBD patients analysed in this association study had reported Jewish ancestry. The lack of an association of this variant with IBD in the Jewish population could reflect the fact that *NLRP7* variants are challenging to genotype because of homology with the nearby gene *NLRP2*, or that environmental components such as commensal bacteria and other factors that act through the *NLRP7* pathway are also likely to play a substantial role in the aetiology of IBD in this population. The potential role of environmental factors is also consistent with the incomplete penetrance of the *NLRP7* variants in families GS13 and GS64.

The *NLRP7* gene encodes the NACHT, Leucine Rich Repeat and Pyrin domain containing 7, protein which is a member of the nucleotide oligomerisation domain (NOD)-like receptor family of proteins. These are a family of pattern recognition receptors which include the known CD risk gene *NOD2* and are sensors of pathogen-associated molecular patterns (PAMPs) involved in the innate immune response, apoptosis and tissue damage. Over recent years there has been increasing evidence for the importance of these receptors in mucosal immunity ^21^ and IBD ^22^ with colonic expression of a number of NLRs, including *NLRP7* being shown to be significantly altered in patients with active IBD ^22^. *NLRP7* expression is induced in response to LPS and IL-1β in peripheral blood mononuclear cells (PBMC) and is detected at high levels in thymus, spleen and bone marrow, suggesting a role in host defence. NLRP3 mediated inflammasome assembly has been demonstrated to be important in both human and mouse models of IBD ^23^ and similarly it is known that the *NLRP7* protein can promote both positive and negative regulation of inflammasome activity ^24^. For example, it is required for bacterial acylated lipoprotein-mediated Caspase-1 activation and maturation of IL-1β and IL-18, but has also been shown to inhibit NLRP3 and Caspase-1-mediated IL-1β release. This contradiction may reflect a role for *NLRP7* in preventing inflammasome formation and IL-1β release in quiescent cells while activating the inflammasome in response to bacterial infection ^25^. In either case, this would be consistent with the association of *NLRP7* mutations with immune-mediated disorders such as IBD. Mutations in *NLRP7* also contribute to the development of hydatidiform mole in abnormal human pregnancies, which may involve inflammation-dependent or independent functions ^26^.

In conclusion, we propose that rare coding variants in *NLRP7* may contribute to the development of IBD. Further work will be required to demonstrate the functional effects of these mutations in the context of intestinal inflammation, and to examine more generally the contribution of aberrant regulation of the inflammasome in the pathogenesis of IBD.

## Author contributions

AO performed experiments and, along with NJP, a significant amount of the analysis and writing the manuscript; KS, NJP, AO and AK performed some specific experiments; YO and AA were involved in sample processing and preparation; KT, RP, BH, JCM & JDS were involved in providing samples from patients and interpretation of clinical data; MAS implemented the exome sequencing pipeline; KdL and JCB provided GWAS meta-analysis data; CGM and NJP conceived, designed and supervised the study, and contributed to manuscript revision.

### ACKNOWLEDGEMENTS

We wish to acknowledge Mr. Philip Tombleson for his assistance with the patient database and the Biomedical Research Centre Genomics Facility at Guy’s and St Thomas’ NHS Foundation Trust and King’s College London. We would also like to thank all the patients and their families who have kindly contributed samples to this study.

